# Emergent particle collection by cyanobacteria through gliding motility and filament buckling

**DOI:** 10.64898/2026.04.28.721293

**Authors:** Rebecca N. Poon, Kelsey Cremin, Alberto Scarampi, Mary Coates, Albane Théry, Orkun S. Soyer

**Affiliations:** School of Life Sciences, The University of Warwick, Coventry, CV4 7AL, United Kingdom; Mathematics Institute, The University of Warwick, Gibbet Hill Campus, Coventry, CV4 7AL, United Kingdom

**Keywords:** Foraging, Cyanobacterial mats, Stromatolites, Emergent behaviours, Microbial aggregates, Plectoneme formation

## Abstract

Cyanobacterial macrostructures such as mats, aggregates, and stromatolites are observed in diverse habitats. These structures incorporate organic and inorganic matter and form microenvironments enabling biochemical transformations. Macrostructures are found to be associated with motile, filamentous cyanobacteria, but how, and to what extent, filament motility can drive macrostructure formation are unclear. To address this question, we study macrostructure formation in a well-characterised freshwater cyanobacterial community dominated by the filamentous cyanobacterium *Fluctiforma draycotensis*. We discover that filament gliding motility leads to the emergence of particle collection behaviour and subsequent aggregate formation at the multi-filament level. This results in aggregate macrostructures comprising a solid core surrounded by an outer layer dominated by entangled cyanobacterial filaments. Developing a novel 3D model of active filament movement, we analyse how filament morphology and mechanical properties affect movement dynamics, particle collection, and filament entanglement. We predict that particle collection requires filaments above a certain length and flexibility. We confirm the resulting prediction of length dependence of particle collection behaviour with shortened filaments of *F. draycotensis* and naturally short *Pseudanabaena sp.* filaments. Together, our results show that filamentous cyanobacteria can use gliding motility to actively engineer their environments through particle collection and macrostructure formation, and that this ability is confined to a part of the filament phase space in terms of length and flexibility. These insights will allow better prediction of macrostructure formation in natural habitats and specific cyanobacterial species, and the engineering of cyanobacterial macrostructures for biotechnological applications.

## Introduction

Cyanobacteria form a diversity of macroscopic structures in aquatic and terrestrial ecosystems. Aggregates and granules can form in aquatic environments, including lakes, open ocean, and bioreactors [1, 2, 6–9], while biofilms and mats are found on rocks or sediments in environments including hot springs and tidal flats [10]. Macrostructures often contain different bacterial species in multiple distinct layers [10–12]. The metabolic activities of resident microorganisms generate gradients of light, oxygen and redox potential, and cycling of key metabolites such as hydrogen and sulfur [7, 8, 13–18]. Therefore, cyanobacterial macrostructures form key habitats enabling microbial transformations of (in)organic matter and are thought to have played a key role in the original oxygenation of the Earth’s atmosphere [10, 13, 19].

Macrostructure morphologies may be smooth or pustular, and may also generate protruding laminated structures known as stromatolites [10, 20]. Cyanobacterial mat sections transferred to the laboratory show vertical and conical growth through accumulation and burial of organics [1, 11, 20–23]. Mats can also incorporate inorganic or nutrient-rich sediment particles, aiding in sediment stabilisation [24, 25] and mineral deposition [26]. Similarly, marine cyanobacterial aggregates involving species of the *Trichodesmium* genus are shown to attach to and incorporate air-borne dust [27, 76, 77]. Cyanobacteria can act as adsorbing, passive surfaces [10, 28, 29], contributing to these macrostructures through adsorption and flocculation. However, other mechanical drivers of particle incorporation and determinants of macrostructure morphology remain unknown.

Environmental macrostructures are often specifically associated with motile, filamentous species of cyanobacteria [2, 10, 27]. Genes associated with gliding motility – particularly those involved in the synthesis and secretion of motility-enabling polysaccharides – are present across nearly all filamentous cyanobacteria [30]. The extracellular polymeric substances (EPS) secreted during gliding motility and broadly known as ‘slime’, can also act as a flocculating agent and cause filaments to attach to each other and to organic substances [28, 29]. Diverse filamentous cyanobacteria form different types of macrostructures in the laboratory [6, 7, 31], but the exact role of gliding motility in the formation dynamics of macrostructures is unclear. Therefore, the cyanobacterial factors influencing macrostructure formation and eventual morphology remain an open question [10, 11, 20] and the ‘engineering’ of cyanobacterial granules for biotechnological applications remains empirical and unpredictable [2, 8].

Here, we studied macrostructure formation in detail, using a motile filamentous cyanobacterium (*Fluctiforma draycotensis*; order Cyanobacteriales based on Genome Taxonomy Database (GTDB)[60, 61]) that reproducibly forms macroscopic granules (‘macrostructures’) in independently initiated cultures and under defined, stable laboratory conditions. This allowed us to track structure formation across macroand microscopic lengthscales, and on timescales from seconds to days. We discovered an emergent behaviour where filament gliding motility generates dynamic collection of solid particles, which are incorporated into granular macrostructures. Combining experiments with a novel physical model of filament propulsion and mechanics, we identified that this behaviour depends on filament length and flexibility, and showed that shortened filaments of *F. draycotensis* do not collect particles to form granular structures. A contrasting short motile filamentous species, *Pseudanabaena sp.* (order Pseudanabaenales [60, 61]), also does not perform particle collection, as predicted by our model. We thus establish a clear role for gliding motility and filament physical properties in dictating macrostructure formation and morphology. These insights and future analyses of our presented model systems will enable better understanding of environmental cyanobacterial macrostructures and their engineering for biotechnologies.

## Results

### Macrostructure formation is driven by gliding motility and particle collection

We studied macrostructure formation first in a microbial community dominated by a motile, filamentous cyanobacterium, *F. draycotensis* (GTDB order Cyanobacteriales; Figure S5) [7] (see *Methods*). This species shows a form of gliding motility characteristic of many filamentous cyanobacterial species, involving surface-attached back-and-forth movement, primarily along the long axis of the filament [32] (Movie S1). The filaments also rotate around their long axis as they move, with the rotation tightly coupled to forward translation [32]. We focused on the repeatable emergence of granular macrostructures in *F. draycotensis* cultures as a model system, then explored another cyanobacterial culture dominated by an alternative motile, filamentous species, *Pseudanabaena sp.* (order Pseudanabaenales). For *F. draycotensis* cultures grown under stable conditions without shaking (see *Methods*), macrostructures formed within 1-2 days of culture initiation and persisted over the lifetime of the culture, which can be up to several months (Figure 1A and *Methods*). Mature granular macrostructures formed under static conditions were highly robust to agitation by swirling or shaking, and comprised an envelope of cyanobacterial filaments surrounding a mm-scale solid yellow core that is enriched in iron compared to the media (Figures 1B and S1). Yellow flocculate iron precipitates were observed to form in non-inoculated media, but they always remained sub-mm and readily dispersed upon minimal agitation (Figure S2). Macrostructures were not observed in cultures containing a mutant, non-motile *F. draycotensis*, or cultures containing only the other community members without *F. draycotensis* (Figure S2). These results suggest that macrostructure formation in this system is driven by the motility of the filamentous cyanobacteria.

**Fig. 1.**
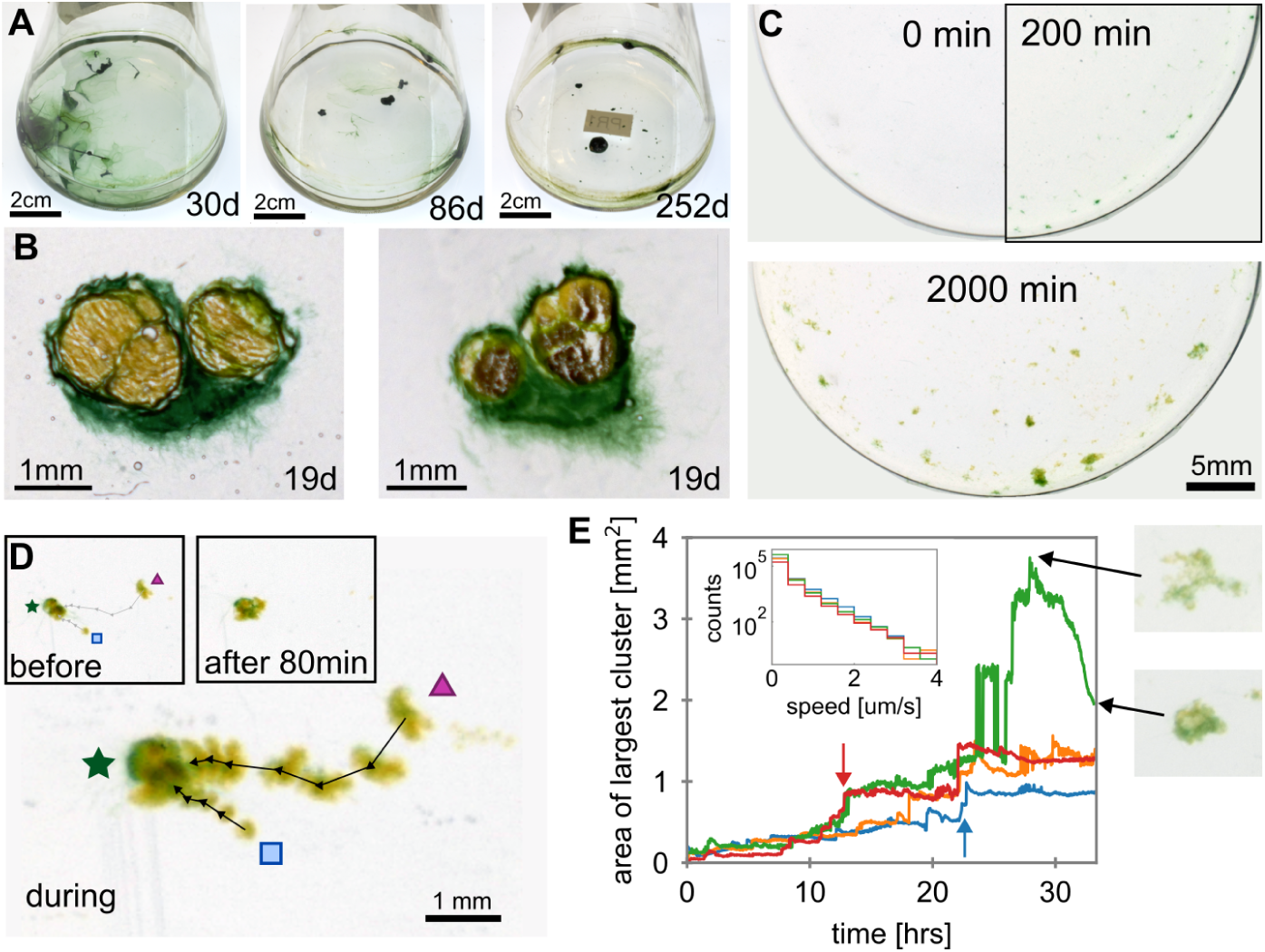
Granule formation at the macroscale. **(A)** Images showing cultures at 30, 86, and 252 days of growth. **(B)** Slices through 19 day old granules, showing green cyanobacteria, and yellow cores containing solid precipitates. **(C)** Images of aggregate formation in a glass beaker, at 0, 200, and 2000 minutes (see Movie S2). **(D)** Time-projected, composite image showing aggregate growth through coalescence of two smaller particles (indicated by triangle and square), with a larger particle (indicated by star). Arrows show the trajectory of the small particles. Insets at top show individual images before (0min) and after (80min) the coalescence event, respectively (see Movie S2 and S3 for example coalescence events). **(E)** Maximum particle size within field of view, plotted against time. Different colours indicate data from separate experiments. Coalescence events cause visible ‘steps’ in the maximum particle size (red and blue arrows). An aggregate ‘tightening’ event is also visible in the data – see still images taken before and after ‘tightening’ event. Inset shows the distribution of aggregates’ speed over the entire video

To understand the formation of these granular macrostructures with embedded particles, we imaged a dilute, homogenised culture in a glass beaker of diameter *≈* 4 cm, over a period of approximately 48 hours (Figure 1C and *Methods*). Over both biological and technical replicate experiments, we reproducibly observed the emergence of highly motile granular structures, which grew to a diameter of 1 mm within 24 hours (Movie S2). Specifically, small yellow particles first appeared at the bottom of the beaker, often surrounded by visible bundles of cyanobacteria filaments. These particles moved with, and were actively collected by, cyanobacterial filaments (Movies S2, S3). Increasingly larger granular structures were formed through coalescence of two or more smaller structures, which appear yellow or green depending on the density of cyanobacteria around the structure (Figure 1D and Movie S3). We quantified the size and speed of macroscopic structures over the course of a 33-hour filming period (Figure 1E and *Methods*). The size increase over this period is mainly due to the macrostructure formation dynamics, as the biomass doubling time in *F. draycotensis* cultures is in the order of one week [7]. We find that structures of 1 mm diameter form after about 20 hours (Figure 1E). Over the course of the entire video we measure speeds of 1-4 *µ*m/s (Figure 1E inset), comparable to measured gliding speeds for individual *F. draycotensis* filaments [32].

In later stages, larger structures were often visibly green due to their dense covering of cyanobacteria (Figure 1A). Macrostructures could also become denser through collected particles and filaments ‘tightening’ (Figure 1E). We found that particle collection by cyanobacteria was not selective for the yellow precipitate particles, and macroscopic polystyrene spheres (430 *µ*m diameter) were also collected if provided, pointing to a physically rather than chemically-mediated process (Movie S4).

### Particle collection is associated with buckling instabilities

We next investigated how particle collection happens mechanistically at the microscopic level, and which specific motility behaviours displayed by *F. draycotensis* are relevant. We took timelapse videos at the microscale, using fluorescence microscopy and microscopic polystyrene beads (48 *µ*m diameter, see *Methods*), and observed the dynamics of bead collection (Movie S5). In a representative example shown in Figure 2A, initially homogenously distributed beads were collected into one of two large (*≈* 100 *µ*m) clusters after 6 hours, incorporating 60% of the beads within the field of view. The time evolution of cluster size distribution at this microscopic length scale is reproducible over experiments initiated on different days using samples from different cultures (Figure 2B).

**Fig. 2.**
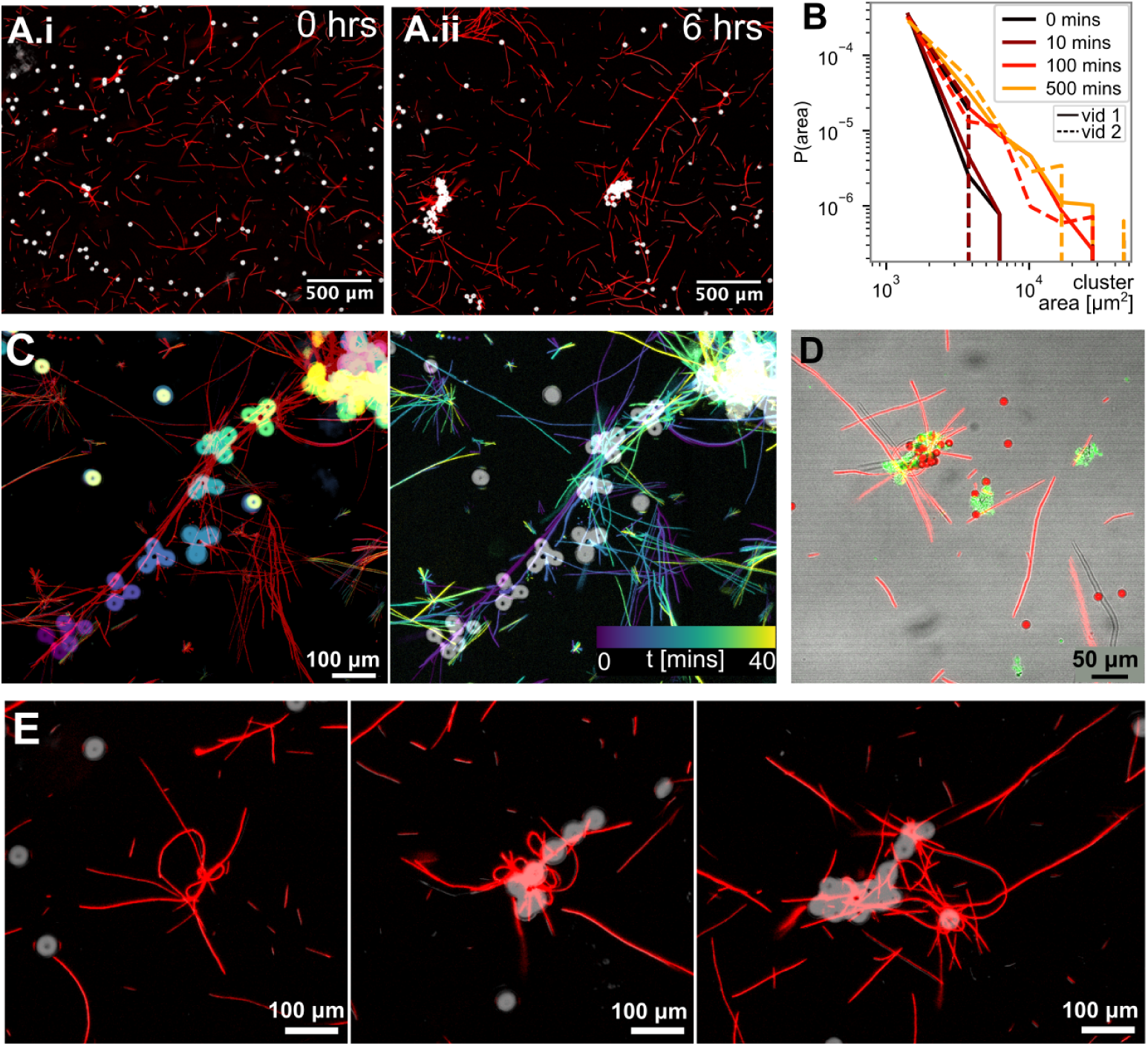
Collection of beads at the microscale. **(A)** Still images of *F. draycotensis* filaments (red) and polystyrene beads (white) at (*i*) the beginning and (*ii*) end of a 6-hour video (see Movie S5). **(B)** The cluster size distribution for the polystyrene beads as a function of time (shown in different colours), from two independent time-lapse experiments (dashed or solid lines). **(C)** Time-projected composite images showing a bundle of filaments and beads transported by their activity. Time is false-coloured from purple (0min) to yellow (40min) and is applied to the beads (left) or filaments (right). **(D)** Example image showing slime concentrated within a bead cluster. The slime is stained with fluorescently labelled lectin (green, RCA-120, Vectorlabs), filaments are light red and beads are dark red. **(E)** Three different examples of entangled filament-filament (left) or filament-bead (middle, right) clusters, featuring long, looped filaments.

We observed bead collection in these microscopic timelapse experiments via at least two specific motility behaviours. First, beads were directly transported upon becoming stuck to filaments, or bundles of filaments, performing ‘normal’ back-andforth gliding motility (Figure 2C). Secondly, buckling and plectoneme formation in a filament were observed to transport and collect beads (Movie S5, S6). Buckling can occur when a gliding filament de-coordinates during a reversal. Rotation-translation coupling can subsequently lead to the filament forming a multiply twisted loop called a plectoneme [32]. We frequently observed that filament clusters (with or without beads) contained filaments that were longer than the population average, and were buckled, sometimes forming loops or plectonemes (Figure 2E). Beads encountered by unbuckled gliding filaments were not always transported, but when transported beads encountered another bead cluster, they generally stayed together. A fluorescent lectin, which is known to stain filamentous-cyanobacteria-associated slime [17, 32, 33], stains these bead aggregates (Figure 2D). We have thus directly observed that slime-mediated filament motility, combined with buckling, and plectoneme formation, drives particle collection and cohesion of bead clusters at the microscopic scale.

### Filament physical properties determine buckling, plectoneme formation and entanglement

To test the hypothesis that buckling and plectoneme formation play an important role in particle collection, we sought a better understanding of how these experimentally observed dynamics relate to the motility and physical properties of the filaments.

We developed a dynamical 3D model for slender filaments that move through local active gliding forces and torques. In this model, each filament is characterised by its centreline position and local orientation, and the filament can bend and twist as it glides on the surface to which it adheres, with its dynamics set by the surrounding viscous fluid (see Figure 3A, *Methods* and *Supporting Information*, *SI*). The essential difference between this model and most previous studies on bending and twisting instabilities of elastic filaments such as DNA [34–36] and worm-like slender bodies [59] is that the filament itself is active and out of equilibrium, and additionally, the forces and torques are applied locally along its centreline and not at its extremities (see *Methods* and *SI*). In other words, we study here elastic instabilities arising upon reversal-initiated movement of two ends of a single active filament against each other, in an otherwise relatively rigid filament. We also resolve the dynamics of the system rather than steady-state configurations: the viscous drag from the surrounding fluid controls the twist and bend propagation along the filament, as discussed in [57].

**Fig. 3.**
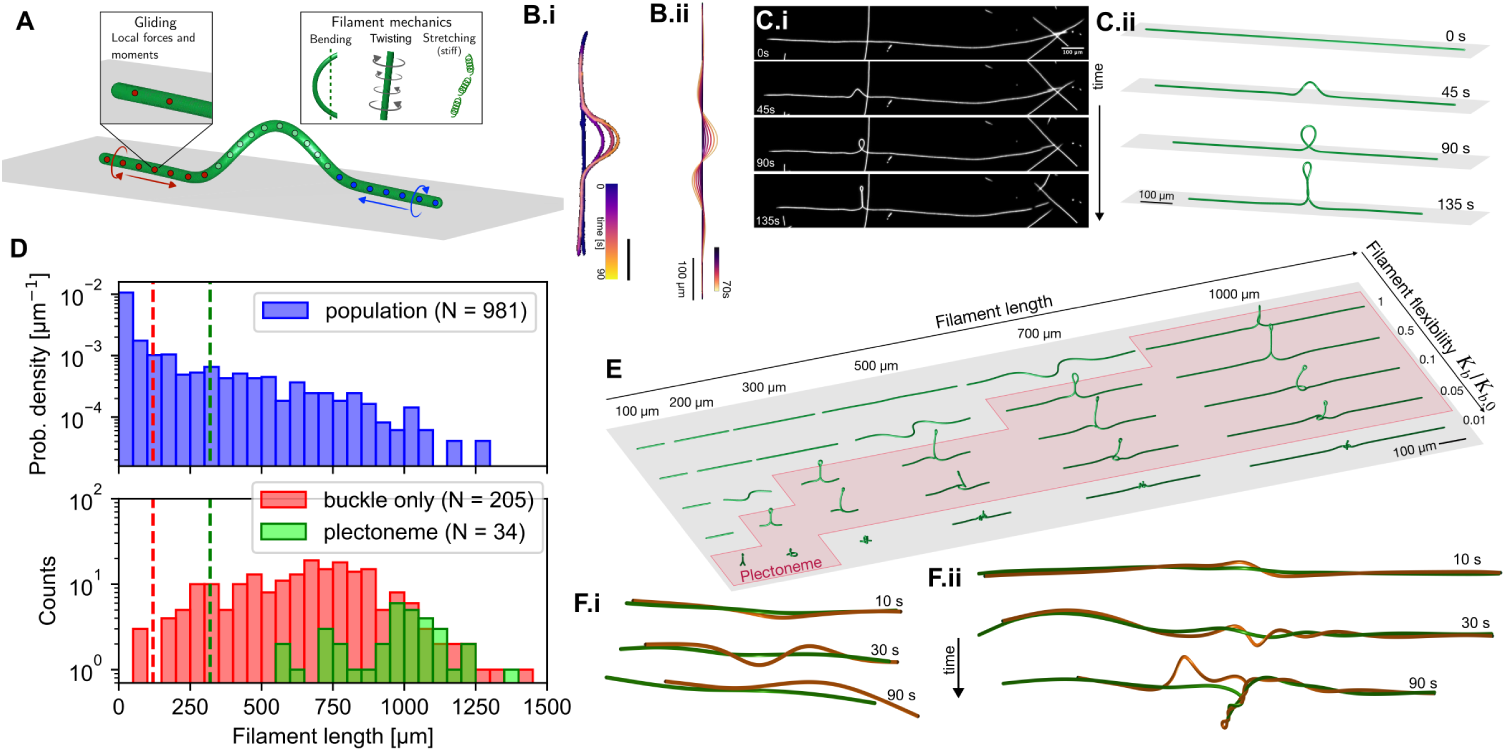
A physical model of cyanobacteria as an active elastic filament. **(A)** Sketch of a simulated cyanobacterial filament. Gliding on the surface is encoded by local forces and moments distributed along the filament length. The filament can bend, twist, and (minimally) stretch, with friction due to surrounding viscous fluid. The coupled rotational and translational directions of the forces are prescribed for a de-coordinated reversal. **(B)** Time-composite images of (*i*) a buckling *F.draycotensis* filament and (*ii*) an exemplar simulated filament. False-colour shows time as indicated. Scale bars 100 *µ*m. **(C)** Images of (*i*) a *F. draycotensis* filament and (*ii*) a simulated filament forming a plectoneme (see also Movie S7). **(D)** Experimental observations of the length distributions of a filament population (blue), and instances of buckling (red) and plectoneme-forming (green) filaments within the population. Dashed vertical lines indicate thresholds for buckling (red) and twisting (green) instabilities predicted by the analytical model (see *Methods* and *SI*). **(E)** A ‘phase space’ for plectoneme formation. Only certain values of length and stiffness (*K*) result in well-behaved plectonemes, as opposed to straight, buckled, or knotted filaments (see also Figure S3 and Movie S8). **(F)** Time series of images from model simulations of pairs of flexible filaments (*K*=0.1), gliding on each other with the orange filament undergoing de-coordinated reversal. (*i*) Short filaments (200 *µ*m) separate and (*ii*) long ones (500 *µ*m) tangle via plectoneme formation (see also Movie 9 and Movies 10-12 for simulated and experimental entanglements, respectively).

We used measured gliding speeds for *F. draycotensis* and bending constants from other motile filamentous cyanobacteria [37, 38] to derive physiologically relevant parameters (Table S1-2), which we then used to simulate buckling and plectoneme formation. These events occur when the leading end of the gliding filament either ‘de-coordinates’ to reverse independently of the trailing end or becomes pinned. The two ends of the filament then produce opposing forces and torques, translating and rotating towards each other (Figure 3A) [32]. For the chosen parameters, the model captures both buckling and plectoneme formation as observed experimentally (Figure 3B, C and Movie S7). The length and flexibility *K* of the filaments determine the type of instability, whilst adhesion and the ratio of twisting to bending modulus (Γ) also control the dynamics and buckled shape (see Figure S3 and Movie S8). We derived analytical expressions for the respective instability thresholds for buckling and twisting instabilities, allowing us to determine critical filament lengths for bending and plectoneme formation of around 120 *µ*m and 320 *µ*m respectively (see *Methods* and *SI*). Both thresholds align well with experimental observations (Figure 3D). The analytical length threshold for twisting instability is slightly lower than the shortest filaments we observed forming plectonemes, which might be because at the critical length, only a small central section of the filament reaches the plectoneme threshold and is not sufficient form a loop (see *SI*).

Model simulations enable us to explore a wide range of parameters for gliding active filaments, resulting in a variety of instability and deformation behaviours, including: absence of buckling/plectonemes; extensive buckling but no plectonemes; plectonemes; and multiple buckling/loops (see Figures 3E, S3, and Movie S8). We found that there exists a limited region of the parameter space, relating to length and stiffness *K*, and to a lesser extent twist-to-bend moduli ratio Γ and surface interaction (see *Methods* and *SI*), in which the experimentally observed single buckling and plectoneme formation occurs (Figures 3E and S3).

Given the observed filament entanglements around collected beads and also the formation of macrostructures in the liquid phase in gently shaken cultures (Figure S4D and *Methods*), we also simulated the interaction of two touching filaments gliding on each other in the absence of a surface. Within the plectoneme-forming parameter region identified above, we found filaments to readily entangle and remain entangled when a de-coordinated reversal occurs, (Figure 3F.i and Movie S9). However, when the interacting filaments are too stiff or short to form plectonemes, then they do not entangle, and only glide past one another and separate (Figure 3F.ii and Movie S9). These simulations qualitatively capture observed entanglement dynamics, and in particular the involvement of plectonemes in initiation of entanglement, which is seen experimentally (Movies S10-12). These model-based results thus show that buckling and plectoneme formation, as well as the associated filament entanglement, are confined to a limited parameter space, requiring filaments with certain physical properties.

### Shorter gliding filaments cannot collect particles nor form granule macrostructures

It is difficult to test all model predictions experimentally. However, filament length is one crucial parameter affecting buckling and plectoneme formation that can be explored. To do so, we took the *F. draycotensis* cultures through a homogenisation and filtering protocol (see *Methods*) to create ‘shortened’ and ‘lengthened’ populations of filaments. The length distributions of the two populations were quantified from small samples taken for microscopy: the maximum filament lengths observed in the short and long populations were 240 *µ*m and 560 *µ*m respectively, with mean lengths of 40 *µ*m and 60 *µ*m respectively. Thus, both short and long populations contained some filaments longer than our estimated critical buckling length of 120 *µ*m, but only the long population contained filaments longer than the plectoneme-forming threshold length of 320 *µ*m (Figure 4A). We then quantified macrostructure formation in these two populations. The shortened population showed gliding motility with similar characteristics to the lengthened population (Figure 4A), but they did not perform particle collection or macrostructure formation over the time window in which the longer population did (Figure 1A and Figure 4B, C).

**Fig. 4.**
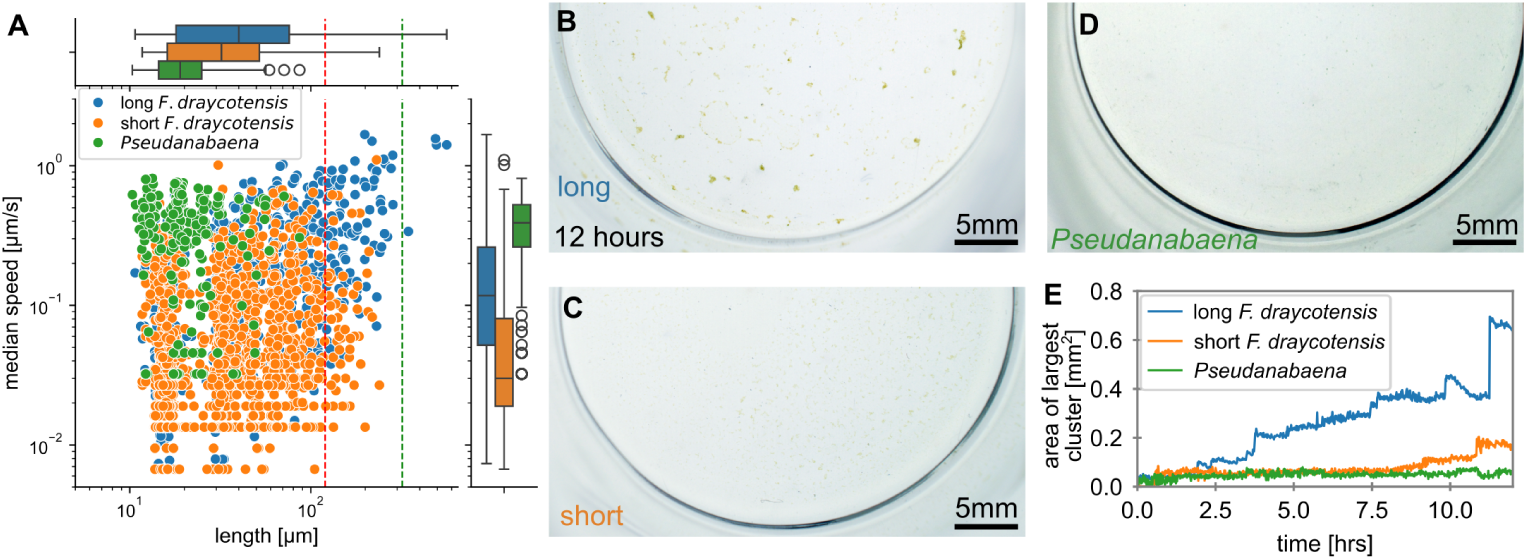
Shorter cyanobacterial filaments do not collect particles. **(A)** Gliding speed and filament length data for three populations of filaments: long (blue) and shortened (orange) *F. draycotensis*, and *Pseudanabaena* (green) (see Movie S13). Dashed vertical lines indicate analytically predicted critical filament lengths for bending (red, 120 *µ*m) and plectoneme formation (green, 320 *µ*m), as in Figure 3D. **(B-D)** Images of the endpoint (12 hours after initiation) of the macro-beaker collection/aggregation assay for the same three populations. **(E)** Time evolution of the largest particle area visible in the beakers for the three populations, coloured as in (A).

To further test the role of length in particle collection, we quantified the motility and macrostructure formation in an additional species of filamentous cyanobacteria: *Pseudanabaena sp.*, which is taxonomically divergent from *F. draycotensis* (see *Methods* and Figure S5), but shows gliding motility with slightly faster speeds than *F. draycotensis* and similar back-and-forth movement (Movie S13). However, they are considerably shorter than *F. draycotensis*, with a mean filament length of 20 *µ*m (Figure 4A). They are therefore predicted by the physical model to be unable to form plectonemes, and indeed, we observed no such instabilities in timelapse videos of *Pseudanabaena* motility, taken over comparable timescales to *F. draycotensis* data in which plectonemes were observed (compare Movies S1 and S13). Using our macroscopic assay of filament behaviour, we observed no particle collection nor aggregate formation by the *Pseudanabaena* filaments (Figure 4D). Multiple days of culture growth under static conditions produced biofilm structures instead of granular macrostructures (Figure S6).

We conclude that, while motile filaments do generally form macrostructures, the nature of these structures depends on their physical properties. Noting that *F. draycotensis* and *Pseudanabaena sp.* belong to deeply divergent cyanobacterial lineages, such primarily physical characterisation may be more significant than other physiological variations in determining the observed diversity of macrostructures formed by different species of filamentous cyanobacteria.

## Discussion

Here we presented and characterised a surprising, emergent behaviour in motile filamentous cyanobacteria: active collection of solid particles from the environment and incorporation of these into persistent macroscopic granular structures. In particular, our results show that the ability of filaments to buckle and form plectonemes promotes filament entanglement and particle collection, thereby leading to the construction of granular macrostructures. The capacity for buckling and plectoneme formation is unlikely to be confined to a single clade; rather, it should arise in any species whose filaments rotate as they translate, and satisfy critical length and flexibility criteria derived here. This insight, as well as the presented modelling and experimental approaches and model systems, will allow development of a predictive framework for understanding the diversity of macrostructures observed in different natural and biotechnological settings and in different filamentous (cyano)bacterial species.

One important conclusion of this study is that filament gliding motility, of the type where filament translation and rotation are coupled, underpins particle collection and dynamical aggregation. We reached this conclusion based on several independent experiments showing an absence of particle collection: (i) in the community featuring a non-motile *F. draycotensis*, and with mostly the same other bacterial species as the community featuring the motile *F. draycotensis*, (ii) in a bacterial community derived from the original *F. draycotensis* community but lacking any cyanobacteria, (iii) in the original community with physically shortened *F. draycotensis*, and (iv) in another cyanobacterial community featuring different bacteria and a naturally shorter, filamentous gliding cyanobacterium, *Pseudanabaena sp*. In addition, timelapse microscopy imaging repeatedly showed clusters of beads being moved by gliding filaments, and thereby being collected into larger clusters. A parsimonious explanation that satisfies all these observations is that particle collection is underpinned by the physical dynamics of gliding and rotating filamentous cyanobacteria, which we show to be linked to specific features of filaments such as their length and elasticity. There may also be involvement of specific molecular factors such as slime, or pili. Type IV pilus machinery, in particular, has been implicated in the gliding motility of non-rotating filamentous cyanobacteria of genus *Nostocales,* and slime has been implicated in the gliding motility of several filamentous cyanobacterial species [30]. We note, however, that both cyanobacteria compared here, *Pseudanabaena sp.* and *F. draycotensis*, encode the type IV pilus machinery together with the hormogonium-associated *hmpD*, *hmpF* and *hmpG*, and hence, the absence of particle collection in only one of them is not explained by absence/presence of these specific proteins. While we cannot account for the possible role of any and all possible molecular mechanisms, we conclude that the presented results are consistent with particle collection being underpinned mainly by the gliding and rotation of long, elastic filaments that can buckle and form plectonemes.

In the current study, we focussed on understanding how particle collection and aggregation formation relate to the physical aspects of gliding motility. We believe that this approach is important to establish a ‘null expectation’ from the physics of gliding, elastic filaments alone. We note that the dynamics of these processes could additionally be influenced by behavioural responses of filamentous cyanobacteria to each other and to external factors, such as responses to light, quorum sensing molecules, or photosynthesis generated radicals. In particular, photo-responses are reported in many gliding filamentous cyanobacteria at the level of individual gliding filaments [3–5, 58]. In the case of species of genus *Trichodesmium*, responses to light are also reported at the aggregate level, and this has been suggested to link to filament reversal dynamics being affected by filament-filament interactions [31]. In the future it will be interesting to scale up the presented model to include multiple interacting filaments, and to systematically test the respective roles of active coordination behaviour for one individual filament (e.g. cell coordination during reversals) and for multiple interacting filaments (e.g. contacts modulating reversal activity), as well as the physical properties (e.g. filament length and flexibility). Such modelling studies can then identify if a ‘purely physical’ model can or cannot generate realistic aggregates, and pinpoint whether additional coordination mechanisms are needed to regulate aggregation. By testing the combination of different physical and biological coordination mechanisms, this would then help to understand better how motility-driven complex behaviours emerge from a physics-biology interface.

While we have described here particle collection and macrostructure formation under laboratory conditions, granule formation and particle attachment has been described before in natural settings such as cyanobacterial mats [10, 25] and marine aggregates, especially of species of the genus *Trichodesmium* [76] but also in other species [78]. For example, the *F. draycotensis* cultures studied here are obtained from a freshwater reservoir, where granular aggregates are commonly observed in the water column, rock pools and on shore rocks (Figure S4). The presented model systems will help us to study these processes under more specific environmental and biological scenarios, such as in the water column or under different flow regimes on surfaces. In the laboratory, we observed *F. draycotensis* macrostructure formation in the bulk liquid phase of either static or gently shaken cultures (Figure S4). However, further systematic studies are needed to quantify granule formation in other settings, such as varied flow conditions or low filament densities. The presented findings and their future extensions will be useful in biotechnology, to help with bioengineering of cyanobacterial granules for water remediation applications [2, 8] using targeted, suitably long and flexible filamentous species and culturing conditions.

These results call for studies of macrostructure formation in conjunction with a stronger emphasis on quantification of filament physical properties. Notably, the presented physical model predictions, regarding the filament length and stiffness required for plectoneme formation and particle collection, were sufficient to explain the difference in collection behaviour in species of two deeply divergent cyanobacterial orders, without invoking lineage-specific biochemistry. Filamentous multicellularity and gliding motility are ancestral traits broadly conserved across cyanobacteria [19, 30], suggesting that the presented biophysical principles in this work may extend to other filamentous cyanobacteria found in environmental mats, biofilms, and stromatolites. Plectoneme formation and macrostructure formation may also be present in other filamentous prokaryotes, such as the sulfide-oxidising cable bacteria (*Desulfobulbaceae*), whose long and flexible filaments form dense sedimentary networks that affect local pH and mineral precipitation profiles [55]. This could affect local concentration of reactive mineral phases, with important consequences for sediment geochemistry.

It is notable that we first observed particle collection with iron-containing precipitates. Iron is a common limiting nutrient for cyanobacteria in many environments where it readily precipitates out of solution. Whilst we found particle collection to be a general capability, extending to any other inert solid particle, it is plausible that micron-sized particles that filaments encounter in nature would include mineral precipitates and nutrient-rich organic matter. A key example of this is seen in bloom-forming marine diazotropch cyanobacteria of the genus *Trichodesmium*, which forms aggregates that can trap iron-rich particles [27, 67, 76, 77]. The detailed dynamics of mineral trapping in *Trichodesmium* is not fully understood, but particles are observed to move along filaments and concentrate in aggregate centres [76]. It is likely that mechanisms other than plectoneme formation are at play in *Trichodesmium*, which usually present as thick filaments and have not been reported to form plectonemes. However, gliding of filaments across each other, as well as aggregation, are observed [31, 76]. Collection of particles into macrostructures can be interlinked with embedding of, or subsequent colonisation by, other bacteria. Indeed, cyanobacterial macrostructures are repeatedly shown to be key habitats for the emergence of interlinked microbial metabolisms [7, 12, 13, 16–18, 66]. Such microbial interactions and overall microbial diversity in granules are shown to be disrupted when macrostructure formation is inhibited in *F. draycotensis* communities [7]. Thus, particle collection and macrostructure formation can have significant downstream functional consequences for microbial communities. We find that motility-driven macrostructure formation in *F. draycotensis* is a highly dynamic process, involving emergent particle collection. Not only is it driven by motile filaments, but the resulting macrostructures can also move, disassemble, and reassemble. Other forms of rapid dynamics are observed in cyanobacterial aggregates of other filamentous species, including *Trichodesmium* [31, 65]. These emergent dynamics result from the interaction of thousands of filaments: such population-level behaviours are widely studied in active matter physics. Such studies have found phase transitions in the patterning of certain filamentous cyanobacteria due to densitydependent nematic ordering [56]; and photoresponsive morphologies in multi-filament bundles [31]. The co-development of models and experiments in the readily-culturable filamentous species like *F. draycotensis* will enable further study of these dynamic, emergent complex behaviours in filamentous active matter.

## Supplementary Information (SI)

This article is accompanied by a supplementary information (SI) text, which includes further methodological details, supplementary figures, and also lists supplementary files and videos associated with the presented work.

## Acknowledgements

We acknowledge Lijiang Song in the Mass Spectrometry Facility, Department of Chemistry at University of Warwick (UoW) for the use of facilities and assistance with ICP-MS, and Marc Walker and Christopher Waldron at the UoW Photoemission Research Technology Platform for their help with the XPS analyses. We thank current and past Soyer group members for discussions and Ben Martynoga, Marco Polin, Roman Stocker, Mohamed Abou Donia, and Emanuele Locatelli for their comments on this work.

## Methods

### Culturing of cyanobacterial community

All cyanobacterial cultures are established from a freshwater source (Draycote Reservoir, UK) and are maintained under stable laboratory conditions as described before [7]. Cultures are kept in the laboratory under regular serial sub-culturing in BG11+ medium (DSMZ medium reference number 1593), with additional vitamins [7]. Subculturing was done approximately monthly, with cultures grown under 12 h/12 h light/dark cycles with white, fluorescent illumination of 14-20 *µ*mol photons m*^−^*^2^ s*^−^*^1^ at room temperature under static conditions. For each passaging step, we performed a 1:50 dilution by transferring 150 µL of resuspended filamentous culture into a final volume of 30 mL of BG11+.

### Granule sectioning

Macrostructures of different ‘age’ and size were removed from cultures using a widebore pipette tip to avoid deformation. Structures were fixed using a glutaraldehydelysine fixation process, adapted from a method for preserving bacterial capsules [39]. Firstly, photogranules were placed in PBS for 5 minutes to remove amines. The granules were then transferred between a series of solutions, all prepared in PBS at room temperature: 50 mM lysine (10 minutes), then 50 mM lysine and 2.5% glutaraldehyde (10 minutes), then 2.5% glutaraldehyde (2 hours), finally followed with 3 transfers into PBS (5 minutes each). Photogranules are snap frozen within a PVP:HMPC hydrogel matrix (2.5% PVP and 7.5% HMPC, g/g) for cryosectioning, based on methods used in [40]. A Bright OTF7000 Cryostat was used to perform cryo-sectioning (chamber: *−*20*^◦^*C, sample head: *−*14*^◦^*C). Samples were sectioned to a thickness of 100*µ*m, and collected onto cover glass slides for imaging.

### Iron analysis of granule cores

A *F. draycotensis* culture was allowed to grow under normal growth conditions described above and regularly checked for visible yellow particles over two days. Emerging, visible particles are collected by pipette under sterile conditions and placed into collection tubes containing high-purity water. These are centrifuged (10,000 *× g*, 10 minutes) to pellet the yellow material, the water is removed and the granules resuspended into 70% ethanol solution for 30 minutes to terminate cell growth. The collection tubes are centrifuged again, and washed three times with 70% ethanol. The material is then centrifuged, and resuspended again in ultra-pure water. The water wash step is repeated twice to remove ethanol. The sample is then placed into a vacuum condenser (Genevac) to remove the water.

### X-ray Photoelectron Spectroscopy (XPS) of granule cores

XPS was performed by attaching a sample of the extracted granule core to electrically-conductive carbon tape, mounted onto a sample bar with a layer of filter paper between the samples and the sample bar to ensure electrical isolation and hence mitigate differential charging, before being loaded into a Kratos Axis Ultra DLD spectrometer which possesses a base pressure below 1 *×* 10*^−^*^10^ mbar. XPS measurements were performed in the main analysis chamber, with the sample being illuminated using a monochromated Al K*α* x-ray source (h*ν* = 1486.7 eV). The measurements were conducted at room temperature and at a take-off angle of 90° with respect to the surface parallel. The core level spectra were recorded using a pass energy of 20 eV (resolution approx. 0.4 eV), from an analysis area of 300 *×* 700 microns. The work function and binding energy scale of the spectrometer were calibrated using the Fermi edge and 3d5/2 peak recorded from a polycrystalline Ag sample prior to the commencement of the experiments. To prevent surface charging the surface was flooded with a beam of low-energy electrons from a charge neutraliser throughout the experiment and this necessitated recalibration of the binding energy scale. To achieve this, the C-C/C-H component of the C 1s spectrum was referenced to 285.0 eV. The data were analysed in the CasaXPS package using Shirley backgrounds and mixed Gaussian-Lorentzian (Voigt) lineshapes. For compositional analysis, the analyser transmission function has been determined using clean metallic foils to determine the detection efficiency across the full binding energy range.

### Inductively coupled plasma mass spectrometry (ICP-MS) analysis of granule cores

Cultures were fractioned into supernatant, cell pellet and yellow granular core, as explained above. Samples of BG11+ media was also included for analysis, to serve as a media reference. All sample fractions were placed in the Genevac and dried until total dryness was achieved. Dried sample was then dissolved in 70% high-purity nitric acid, diluted into water (1:20) and then filtered through 0.22*µ*m PES filters, and analysed on the ICP-MS. A SPS4 auto-sampler Agilent 7900 was used for ICP-MS. The following equipment conditions were set: spray chamber temperature: 15*^◦^*C, plasma gas flow rate: 15 L min-1, carrier gas flow rate: 0.9 L min*^−^*^1^, nebuliser gas flow rate: 1.05 L min*^−^*^1^, auxiliary gas flow rate: 0.9 L min-1, and helium mode gas flow rate: 4.0 mL min*^−^*^1^. The forward power was 1550 watts, with a flow rate of 0.1 rps. Detector analog HV 2152V, pulse HV 956V, with a discriminator voltage of 5 mV and a 4.3 mL helium gas flow rate. Prior to the sequence, the instrument was auto-tuned, and during the analysis, the P/A was modified using calibration standards.

### Shaken cultures

For the ‘swirling’ flask shown in Figure S4D, cultures were photographed after 16 days of growth. The culture was grown in an Algem 1.4 (ALGae Environment Modelling lab scale photobioreactor), by Algenuity, at a rotation rate of 80 rpm. The flask was illuminated from beneath with LEDs at *≈* 60 *µ*mol photons m*^−^*^2^ s*^−^*^1^, on a 12hr/12hr light/dark cycle. The culture was set up using a 1:50 dilution (8ml culture in total 400 ml BG11+ media).

### Macroscopic aggregate formation experiments

The process of aggregate formation was recorded on a macroscopic scale in 50mL glass beakers of approximately 37 mm diameter. The beakers were inoculated with 0.4 mL of a mature culture sample, aged between 35 and 70 days, and diluted 1 in 50 into 19.6 mL of fresh BG11+ media to give a total fluid volume of 20 mL. As the microbial communities found in culture flasks of this age have usually formed persistent, cohesive granules, we homogenise the inoculant to break up these structures before setting up the beaker experiments. We do this by pipetting approximately 1 mL of sample up and down in an Eppendorf tube. We then dilute the homogenised inoculant to an optical density of 0.4 compared to a media blank (absorbance at 750 nm, measured using a SPECTRONIC 200 Spectrophotometer, Thermo Fisher Scientific).

The experiment is set up by placing the beakers containing only the fresh media under a DSLR camera (Nikon) held on a post at sufficient distance to allow the entire base of the beaker to be focused in the imaging plane. The beakers are lit from below using a sheet of LED lights, diffused through multiple layers of white diffusion paper (Lee Filters, 216 White Diffusion Lighting Gel) to give a brightness of 36*µ*mol photons m*^−^*^2^ s*^−^*^1^ at the base of the beakers. The camera is focused by hand at the beginning of the experiment, with adjustments made during the imaging time as necessary. Once the initial focus is set, the sample is added to the media in the beaker, and the beaker is covered with a plastic petri dish lid to reduce evaporation. Images are then taken once every minute for up to 2000 minutes (33.3 hours).

The macroscopic collection assay in Figure S2 was performed in the same glass beakers as all other ‘macroscopic aggregate formation’ experiments. For Figure S2A, we used higher initial densities of culture, and rotated the beakers gently at 150rpm (using a Heidolph Unimax 1010 rotary shaker) to physically concentrate any solids to the centre of the beaker via fluid flow. This allowed easy visibility of either active collection (in the case of the motile culture) or precipitation (for the other two cases) after 5 hours. Additional agitation was then performed by manually swirling and tapping the beaker. For Figure S2B, we used a derived bacterial community lacking *F. draycotensis*, grown on BG11+ with added 0.8% (*≈* 44mM) glucose. As this ‘no-cyanobacteria’ community soon becomes optically dense, black polystyrene beads (Thermo Scientific Black ChromoSpheres Polymer Microspheres, 284*µ*m diameter) are added to the beakers to allow particle collection to be easily visualised.

### Macroscopic aggregate formation image analysis

Images of macroscopic aggregate formation within glass beakers are segmented and analysed to measure particle size and speed. Segmentation is performed on the blue channel of the RGB images, which has the largest contrast between the background and the aggregates. The images are automatically thresholded to segment the particles from the background, then particle positions and areas are measured using the Analyze Particles function in ImageJ. The particles are then tracked using the Trackpy module in Python. This code is available at https://github.com/OSS-Lab/Cyanobacteria_particle_collation.

### Microscopy imaging of filaments and beads

Particle collection at the microscale was imaged using an Olympus IX83 inverted microscope with a Photometrics CoolSnap-HQ2 camera. Time-lapse videos were taken in brightfield and fluorescence imaging. Chlorophyll autofluorescence of the filaments was imaged using a CoolLED pE-300 illumination system and a Chroma HcRed1 filter cube (Excitation filter: *≈*550-600 nm, Dichroic mirror: Q610lp, Emission filter: HQ640/50m, *≈*615-665 nm). We used time intervals of 10s-60s and an exposure time of 100ms. Exampler timelapse images are available at https://zenodo.org/uploads/18680430.

Experiments were performed in a 6 cm glass petri dish base, with a plastic petri dish used as a lid. 12mL of BG11+ media was added to the dish, along with a small quantity of 48*µ*m diameter black polystyrene microspheres (ChromoSphere Dry Dyed Polymer Particles, Black, BK050). Between 0.2-0.4 mL of homogenised mature sample of the cyanobacteria was then added and imaging was initiated immediately afterwards.

### Lectin staining

Following one of microscale collection experiments described above, we stained the sample with a fluorescein-tagged lectin stain (RCA-120, Vectorlabs). This lectin agglutinates sugars including galactose [33], which is known to be a major component of the slime produced by *F. draycotensis* [7]. Images of the stained sample were taken using a Zeiss 880 confocal microscope. The RCA-120 lectin was imaged using a 488 nm argon laser, with a PMT emission range of 510-550 nm (green channel), whilst the autofluorescence from the filaments and beads was imaged using a 633 nm HeNe and an emission range on the PMT of 638-700 nm. Imaging was performed with a pinhole diameter of 45.96*µ*m, and a MBD 488/561/633 beamsplitter was used to allow imaging across both channels simultaneously

#### Creation of shortened *F. draycotensis* populations

Populations of *F. draycotensis* with shortened filaments were created by homogenizing *≈* 1.5 mL mature culture, then allowing it to settle for *≈* 30 minutes in a 2 mL Eppendorf tube, resulting in a loose aggregate. The ‘supernatant’ of this aggregate was extracted and filtered through a 40*µ*m mesh (Greiner Bio-One EASYstrainer Cell Strainer, Small) to give the ‘short population’. The loosely aggregated filaments were resuspended and homogenised by pipetting up and down in order to give the ‘long’ population. The length distribution and motility of the two populations were confirmed by direct measurement of microscopy images of small samples, giving mean lengths of 40 *µ*m and 60 *µ*m, and maximum lengths of 240 *µ*m and 560 *µ*m, for the short and long populations respectively (Figure 4A). For inoculating the beaker experiments (Figure 4B-D), the densities of the two populations were matched by optical density measurements.

### Length and speed distributions of filament populations

We used time-lapse microscopy images of filaments gliding on a glass surface to measure length and speed distributions of various populations of filaments. Timelapse images are available at https://zenodo.org/uploads/18680430. Samples were prepared on glass slides with an imaging spacer (Thermo Fisher 25*µ*L Gene frame) and sealed with a plastic cover slip, and imaged on an Olympus IX83 inverted microscope in the HcRed fluorescence channel. The filament density was sufficiently high that any frame of the video contains a significant fraction of filaments that overlap with one another. We therefore implemented an automated segmentation routine to extract the filaments, based on the method outlined in [41]. This code is available at https://github.com/OSS-Lab/Cyanobacteria_particle_collation. The filament trajectories were then linked in Python using the Trackpy module in order to calculate filament speeds.

### Phylogenomic analysis

A phylogenomic tree was constructed from 97 genomes (Figure S5, Supplementary File S1): genomes obtained from laboratory enrichments experiments, including a *F. draycotensis* [7, 42], all GTDB species representatives of the family JAAUUE01 to which *F. draycotensis* belongs, four genomes each from the families Chamaesiphonaceae and Coleofasciculaceae, 1-5 genomes from each of the 19 orders of Strunecky et al. [63], as represented in GTDB release R232 (selected by genus, spread across families, preferring isolate genomes with *≥*90% completeness and *≤*5% contamination), and three non-photosynthetic Cyanobacteriota (Sericytochromatia and Vampirovibrionia) as an outgroup. For reference genomes without an existing gene call, proteins were predicted with Prodigal [74]. The 71 bacterial single-copy core genes of anvi’o (Bacteria 71) [43] were identified with `hmmsearch` (HMMER v3) [73] using the profiles’ gathering thresholds, keeping the best hit per genome. Each gene was aligned with MAFFT L-INS-i [72] and trimmed with trimAl v1.4 [44] (gap threshold 0.5). Genes present in fewer than 85% of genomes were discarded, and the remaining 67 genes were concatenated into a supermatrix of 12,293 amino-acid positions. For two genomes described in [42] (*Pseudanabaena* sp. OSS1 Bin 161 and *Calothrix* sp. NONANO MAG 00047), only the six ribosomal proteins (L1–L6) of the original analysis were available, and their remaining positions were coded as missing data. The maximum-likelihood tree was inferred with IQ-TREE 3 [68] from the partitioned supermatrix, with substitution models and partition merging selected by ModelFinder [71] under the Bayesian information criterion. Branch support was assessed with 100 non-parametric bootstrap replicates under the selected partition scheme, 1,000 ultrafast bootstrap replicates [62] and the SH-aLRT test [46]. Gene and site concordance factors [69] were computed against 67 single-gene maximum-likelihood trees. The tree was rooted on the non-photosynthetic outgroup (Sericytochromatia and Vampirovibrionia). To test robustness to model choice, the tree was re-estimated under the site-heterogeneous LG+C20+F+G model using the PMSF approximation [70], and the resulting tree differed from the presented one at three branches, all with bootstrap support below 65%.

The tree was drawn with Biopython [75] and Matplotlib. Taxonomic labels shown on the tree follow GTDB release R232 [60, 61]. Additionally, the order of each genome under Strunecky et al. [63] is given in Table S2 and Supplementary File S1. The morphology shown on the tree (filamentous or unicellular) was taken from published strain descriptions, the morphological section reported by Yang et al. [64] or the genus description. The habitat information shown on the tree is taken from the original strain description or the isolation source of the NCBI BioSample record; both were left blank where no information was available.

Electronic version of the tree, along with associated sequence files and figure generation code are available at https://github.com/OSS-Lab/Cyanobacteria_particle_collation.

### Filament modelling and simulation methods

#### Filament model

We developed a 3D physical model of slender, elastic filaments undergoing active motion to describe the gliding motility, buckling, and plectoneme formation of the filamentous cyanobacteria. The model code is available at https://github.com/OSS-Lab/Cyanobacteria_particle_collation. This model builds on an existing framework that combines the classical equilibrium theory for passive elastic filaments—formulated through Kirchhoff rod models for systems such as DNA [34–36]— with immersed-boundary methods for describing filament dynamics in a viscous fluid [48–51]. We extend this approach to include the local active forces generated by gliding motility, and interactions with the external environment. Importantly, the model is agnostic about the detailed mechanism responsible for generating the gliding motility (whether type IV pili, slime extrusion, or another mechanism [32]), and only accounts for the minimal experimentally observed elements of *F. draycotensis* motility: the propulsion stems from individual cells, can act either forwards or backwards, and leads to both translation (from active forces) and rotation (from moments). We model both unidirectional motion, where all the cells are synchronised, and decoordinated reversals, where the two ends of the filament move towards each other. We approximate the medium in which the cyanobacteria move as a viscous fluid above a solid surface, which causes friction and sets the dynamics of filament motion. Within this formalism, any filament deviating from its relaxed, straight configuration experiences elastic forces and torques that oppose bending and twisting, and these are transmitted to the viscous fluid as hydrodynamic singularities. The filament then moves and reorients according to the local active flow field [48–51].

#### Filament description

The configuration of the filament can be described by the centreline position **X**(*s*) of the filament at a given point *s* and its local orientation, given by the local material frame, an orthonormal director basis [**d**_1_(*s*), **d**_2_(*s*), **d**_3_(*s*)]. In the classical Kirchhoff framework, **d**_3_(*s*) is the tangent to the curve **X**(*s*), and *d*_1_ and *d*_2_ point normal to the curve **X**(*s*). The twist vector ***κ***(*s*) is then defined from the derivatives of the director basis with respect to the arclength, *∂_s_***d***_i_* = ***κ*** *×* **d***_i_*, where ***κ*** = *κ*_1_**d**_1_ + *κ*_2_**d**_2_ + Ω**d**_3_.

The elements of the twist vector are known as filament curvatures, and the curvature in the third direction Ω is referred to as the twist density. In our dynamic simulations, because the filament moves with the local flow, as in [49], we allow the material frame to deviate from the tangent and normal vector triad as it rotates with the flow. We also allow for some weak extension of the filament, but impose a large stretch modulus *b*_3_ and shear force constants *b*_1_ and *b*_2_.

#### Bending and twisting forces and moments

With these formulations in place, we can write down the equation of state describing the elastic forces (*F*) and torques (i.e., moments, *M*) on a point *s* on the filament at time *t* for a given filament configuration as;

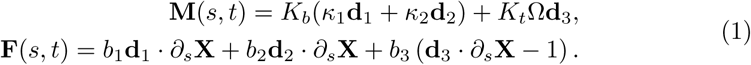

where we introduced the twist coefficient, *K_t_*, and the bending coefficient, *K_b_*, describing the resistance of the filament to twisting and bending, respectively, and *b*_3_ and *b*_1_, *b*_2_ describe its resistance to stretch and shear, with *b_i_ > K_b_, K_t_*.

To characterise the filament mechanics, we introduce the relative bending stiffness *K* = *K_b_/K_B,_*_0_ relative to the mean experimental value from several cyanobacteria filaments *K_B,_*_0_ [37, 38], and the ratio of twist-to-bending moduli, Γ = *K_t_/K_b_*.

#### Active gliding

To model active gliding motility, we incorporate the active forces and torques from each cell of the filament. While the exact underlying mechanism for the gliding is not well understood, we know that in *F. draycotensis* filaments - each cell translates and rotates as it moves [32]. As a minimal model for gliding, we consider that at each point, the filament experiences local forces and moments along the local tangent vector **t**(*s*) with densities **f**_glide_ and **n**_glide_, respectively.

We impose that the gliding only occurs when the filament is in contact with a solid surface or another filament. We use the function *δ*_cont_(*s, t*) which is 1 if the distance of the filament point *s* to the surface or to the nearest filament (*x*(*s*)) is less than *δ*_adh_, 0 otherwise.

The direction of gliding is *α*(*s*) *∈ {−*1, 1*}*. For a forward-gliding filament, *α*(*s*) = 1 everywhere. For a reversal, we impose *α*(*s*) = 1 when *s ∈* [0*, L_f_ /*2] and *α*(*s*) = *−*1 when *s ∈* [*L_f_ /*2*, L_f_*], meaning that the front half of the filament reverses.

Overall, the forces and moments along the local tangent vector **t**(*s*) are described as;

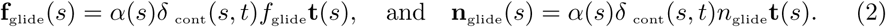

#### Surface interaction

We also impose a steric repulsion between the surface and the filament. The steric interactions are active where the centreline of the filament is touching the surface, *x <* 0, are directed normal to the surface, and increase linearly with *x*, **f**_ste_(*s, t*) = *c*_ste_*δ_x<_*_0_*|x*(*s, t*)*|***ê***_x_*.

Finally, we consider close-range adhesion between the surface and the filament, with a typical strength *c*_adh_, the effect of which we explore in simulations. It is active on portions of the filament that are in the adhesion region *x ∈* [*δ*_adh_ */*2*, δ*_adh_], with 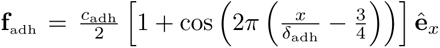, so that the force is oriented perpendicular to the surface, cancels at the surface and top of the adhesion layer, and reaches a maximum *c*_adh_ in the middle of the layer.

The steric interaction strength is taken to be very large compared to the active and adhesion forces, which are assumed to be of the same order of magnitude, *c*_ste_ *≫ c*_adh_ (see SI).

#### Force balance

With the above definitions in place, we can write the force balance on a point *s* on the filament at time *t*. To do so, we use a coupled system of equations where the elastic forces and torques generated by the bending and twisting of the filament are transmitted to the viscous fluid. The rod experiences the elastic forces and torques from eq. (1), active force and torque densities **f**_glide_, **n**_glide_ from gliding, viscous forces and torques from the fluid **f***_v_*, **n***_v_*, and steric interactions with the surface **f**_ste_. The force and moment balance is;

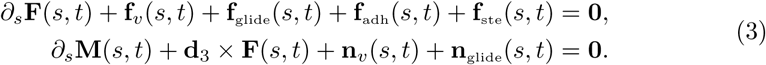

#### Surrounding fluid

Our simulations for the elastic rod in a viscous fluid relies on the immersed boundary method for a slender filament [48–51], which describes the interaction between the active elastic filaments and the surrounding viscous fluid. In this setup, the elastic and gliding forces and torques are directly transmitted to the fluid according to eq. (3), and they generate an active flow around the filament. The filament in turn moves and deforms with the local fluid velocity.

The surrounding viscous fluid verifies the incompressible Stokes equations for the fluid velocity **u** and pressure **p**,

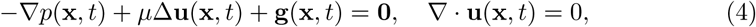

with **g** denoting the distribution of forces per unit volume imposed on the fluid.

As the propulsion, elastic, and adhesion forces on the filament are all transmitted to the fluid as point forces and torques, they generate a local force per unit volume distribution, **g**(**x***, t*). To avoid singularities in the flow velocity, we use a regularisation kernel *ψ_ɛ_* such that the forces are not applied on singular points of the filament but over an area of typical size *ɛ*.

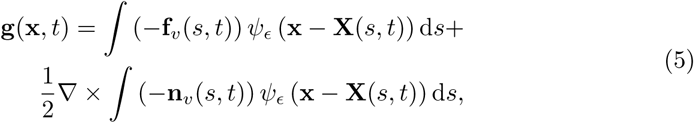

with 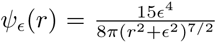. We then compute the local velocity of the fluid given this distribution of local forces according to eq. (4) as the sum of regularised Stokeslet (left integral) and rotlet (right integral) singularities.

In addition, we enforce the no-slip boundary condition at the solid surface **u** ((*−δ, y, z*)*, t*) = **0** by using regularised image singularities for the Stokeslets and rotlets [52–54]. That is, we ensure that the fluid at the solid surface has zero velocity by adding an adequate combination of local flow singularities at the mirror points from the active filament, with their exact expression given in [54].

#### Filament dynamics

The filament then moves and rotates with the local fluid velocity, as

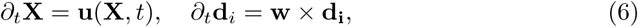

with 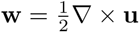 the local flow vorticity.

The numerical implementation for the resulting system follows [49]. We discretise the filament in sections of size d*s* = 1*µ*m, which is of the order of the filament radius. To compute the derivatives of the forces and moments along the arclength, we compute them at midpoints of the filament and use auxiliary reference frames at these midpoints. We then use a forward Euler algorithm to solve the coupled system of equations. In all our simulations, we use parameter values from the literature on filamentous bacteria when available [32, 37, 38], and infer physically consistent values for the other parameters. The parameters used in the simulations are given in Table S1 for physical properties and in Table S2 for numerical implementation, and their influence on simulation outcomes are discussed in the *SI*.

#### Two filament interactions

We extended these simulations to explore the buckling and twisting dynamics of two filaments moving against each other. In the two filament system, gliding along the filament direction occurs for points that are within the contact region of the other filament (instead of the surface) and generate an equal and opposite force on the other filament. We also account for steric interactions between the filaments. We study in particular their entanglement when a reversal occurs in the middle of one of the filaments, as seen experimentally. We find that the plectoneme formation regime (long, flexible filaments) in the two-filament system leads to entanglement, as depicted in Figure 3F.ii. Shorter filaments, which wouldn’t form plectonemes on the surface, instead tend to glide past one another and separate even when a reversal occurs (Movie S9).

#### Simulation parameters

In all our simulations, we use parameter values from the literature on filamentous bacteria when available [32, 37, 38], and infer physically consistent values for the other parameters (see Table S1).

For the motility, we use the experimentally measured gliding force density *f*_glide_ = 1 nN.*µ*m*^−^*^1^ [38]. We assume that bacteria glide over a film of thickness of 1*µ*m above the solid surface, and set the medium viscosity *µ* to match the experimental gliding speed of the filament *v*_glide_ *≈* 1*µ*m*.s^−^*^1^ for a 200*µ*m filament based on *F.draycotensis* filament speeds measured here and previously [32]. We then chose a torque density so that the rotation rate matches its measured experimental value *ω*_glide_*/v*_glide_ *≈* 0.87*µ*m*^−^*^1^ [32]. In order to keep our model minimal and independent of the details of the gliding mechanism, we neglect any non-linear effects in the surrounding fluid, including slip conditions or non-Newtonian fluid rheology that might arise from the slime. We instead keep the fluid viscosity as an effective parameter to set the gliding dynamics. In particular, these values fall well into the viscous flow regime with a Reynolds number Re *≈* 5 *×* 10*^−^*^12^.

The bending of filamentous cyanobacteria has been studied experimentally [37, 38], but we couldn’t find in the literature values for the twisting modulus of filamentous bacteria, and assumed a unit twist-to-bend ratio Γ, but also explored other values in simulations. We take the adhesion force to be stronger than the gliding forces, *c*_adh_ = *f*_glide_. For smaller values of this coefficient, buckling leads to a detachment from the surface, contradicting experimental observations (see Figure 3E and S3). The numerical parameters are chosen for the simulations to be stable, relying on previous implementations of elasto-hydrodynamic simulations [51].

## Declarations

### Funding

This project is funded by the Gordon and Betty Moore Foundation (grant 9200: https://doi.org/10.37807/GBMF9200). We acknowledge the use of computational facilities provided by the Scientific Computing Research Technology Platform (SCRTP) at the University of Warwick.

### Conflict of interest

The authors declare there are no conflicts of interest relating to this work.

### Data and code availability

Data, model and analysis code are available at the following dedicated GitHub and Zenodo repositories: https://github.com/OSS-Lab/Cyanobacteria_particle_collation https://zenodo.org/uploads/18680430

### Author contributions

RP, AT, and OSS conceptualised and developed the methodology of the study. RP, MC, KC, and AS performed experiments and analyses, while AT developed models and ran simulations. RP, AT, and OSS performed formal analysis and visualisation of the data. RP, AT, and OSS wrote the manuscript and all authors contributed to its review and editing. OSS obtained funding for the study, and OSS and MC were responsible for project administration.

## Notes

https://github.com/OSS-Lab/Cyanobacteria_particle_collation

### Competing Interest Statement

The authors have declared no competing interest.

### Summary of Updates

Presentation of results, expansion of discussion, and clarification of methods.

