## Supplementary information for "Emergent particle collection by cyanobacteria through gliding motility and filament buckling"

**This PDF file includes:**

Supporting information (SI) text Tables S1 to S3 Additional supplementary files and their legends; S1 to S13 Figures S1 to S6 SI References

#### Supporting Information

##### Scalings for buckling and plectonemes thresholds

Here, we use scaling laws to derive analytical expressions for length thresholds relating to buckling and plectoneme formation. Note that this system is different from most problems describing the twisting or compression of an elastic rod, in particular for plectoneme formation in DNA [1–3] or in helically shaped, motile bacteria *Spiroplasma* [4]. Those cases involve forces and moments imposed at the extremities of the rod, which lead to constant load and twist densities along the rod until buckling [5, 6]. Here, by contrast, the twisting and compression are produced by local active torques and forces, and the twist density and axial load increase linearly along the filament.

***Twist instability***. Let us first consider the twisting of the rod when local forces at its two ends cause these to rotate in opposite directions and translate against each other, as seen experimentally [7] (see also Movies S10-12). We consider twisting before any bending, and denote by $\Omega$ the twisting density, and $\phi$ the local twist angle ($\phi(s) = 0$ initially). From the constitutive equations of the rod and the force balance, we obtain the following twist equation:

$$K_t \partial_s \Omega = -n_{\text{glide}}(s) - m_{v,||}(s,t) \tag{1}$$

Using resistive-force theory [8], we can write the local viscous torque resisting twist as $m_{v,||}(s,t) = \xi_r \partial_t \phi$, with the drag coefficient $\xi_r \approx 4\pi\mu r^2$. Writing the constant gliding torque on the two counterrotating halves of the filament $n_{\text{glide}}(s) = n_{\text{glide}}\text{sign}(s)$, we obtain a diffusion equation for the twist along the filament [9]:

$$\frac{K_t}{\xi_r}\partial_{ss}\phi = \partial_t \phi - \frac{1}{\xi_r} n_{\text{glide}}\text{sign}(s). \tag{2}$$

We can solve this with initial condition $\phi = 0$ and free ends of the filament ($\partial_s \phi|_{\pm L/2} = 0$) and in particular obtain the steady-state twist density at equilibrium $\Omega_{eq}$, its steady maximum value at the rod midpoint $\Omega_m$

and and the timescale for twist propagation over the rod $\tau_t$:

$$\Omega_{eq} = \frac{n_{\text{glide}}}{K_t}\text{sign}(s)\left(|s| - \frac{L}{2}\right), \quad \Omega_m = \frac{n_{\text{glide}}L}{2K_t}, \quad \tau_t = \frac{L^2\xi_r}{4K_t\pi^2}. \tag{3}$$

In this active filament system, the critical twist density for buckling is not uniformly reached throughout the entire system, and we find that twist accumulates in the central region of the filament and decays linearly to reach zero at the endpoints on both sides. We still leverage the classical results for a rod twisted at the end to obtain scalings for the buckling of an active filament [3]. In particular, we assume that a good approximation for the buckling threshold is when the midpoint reaches the critical twist density $\Omega_{c,t} = \frac{2\pi}{L\Gamma}$ for a free filament, with the previously defined twist-to-bend moduli ratio $\Gamma = K_t/K_b$. Note that we expect this assumption to underestimate the filament length needed to form a plectoneme, which can only occur if the middle section that has reached the twist instability threshold is long enoufgh.

This yields a critical length for which the filament is unstable in twist (when $\Omega_m = \Omega_{c,t}$), $L_{c,t} = \left(\frac{4\pi}{n_{\text{glide}}}K_b\right)^{1/2}$. This expression yields $\approx 320\mu m$ for parameters listed in Table S1, based on measurements from *F. draycotensis* and similar cyanobacteria. Note that this expression for the critical twisting length does not depend on the twisting modulus $K_t$, but the timescale associated with the twist build-up $\tau_t$ scales as $K_t^{-1}$.

***Bending instability.*** We next consider the filament under active compression from a force density $f_{\text{glide}}(s) = -f_{\text{glide}}\text{sgn}(s)$ applied on both ends, but without any rotation ($n_{\text{glide}} = 0$). We assume that the filament is inextensible ($b_i \gg K_b, K_t$), so the forces build up instantly and are transmitted along the rod. The local compression is stronger at the centre of the rod, zero at the free ends, and increases linearly. This setup is similar to self-load problems in Euler buckling.

The axial load on the filament is compressive,

$$N(s) = f_{\text{glide}}L\left(1 - \frac{2|s|}{L}\right). \tag{4}$$

To compute the threshold for buckling, we write the Euler-Bernoulli beam equation for the linear, small-amplitude radial deflection from the straight filament, which we denote $w(s)$ as in [10];

$$K_b\frac{\mathrm{d}^4w}{\mathrm{d}s^4} + \frac{\mathrm{d}}{\mathrm{d}s}\left[N(s)\frac{\mathrm{d}w}{\mathrm{d}s}\right] = 0. \tag{5}$$

The dimensionless length controlling the system is:

$$l_0 = \left(\frac{K_b}{2f_{\text{glide}}}\right)^{1/3}, \tag{6}$$

therefore, the critical buckling length, $L_c$, will scale with $l_0$, $L_c = Cl_0$. We can rescale the problem with a new variable $\xi = l_0^{-1}\left(x - \frac{L}{2}\right) \in [-L/(2l_0), 0]$. This results in an Airy equation for the slope of the local deflection angle $\theta = \mathrm{d}w/\mathrm{d}s$;

$$\frac{\mathrm{d}^2\theta}{\mathrm{d}\xi^2} - \xi\theta = 0. \tag{7}$$

The ends of the filament are free, so at $s = \pm L/2$ (that is at $\xi = 0$), $\mathrm{d}\theta/\mathrm{d}\xi = 0$ (no bending moment). We also impose the symmetry at $s = 0$, which sets $\theta = 0$ at $\xi = -L/(2l_0)$. The solution can be written with Airy functions, $\theta = c_1\text{Ai}(\xi) + c_2\text{Bi}(\xi)$, and the $\xi = 0$ condition yields $c_2 = c_1/\sqrt{3}$. The first buckling instability happens for the first mode where $\text{Ai}(-L_c/(2l_0)) + 1/\sqrt{3}\text{Bi}(-L_c/(2l_0)) = 0$ has a non trivial solution. Numerically, this yields a constant $C \approx 4$, and a minimal length for buckling $L_c \approx 4\left(\frac{K_b}{2f_{\text{glide}}}\right)^{1/3}$.

We can also consider the buckling of a filament of length $L$ when it gets pinned on the head, while gliding forward. In this "pinned by the head" scenario, which is experimentally observed (see Movie S1), the load and characteristic scales are:

$$N_p(s) = f_{\text{glide}}L\left(1 - |s|/L\right), \qquad \text{and} \quad l_{p,0} = \left(\frac{K_b}{f_{\text{glide}}}\right)^{1/3}. \tag{8}$$

We use the rescaled variable $\xi_p = l_0^{-1}(x - L)$. The equation for the local deflection angle $\theta = w'(s)$ is the same, $\frac{\mathrm{d}^2\theta}{\mathrm{d}\xi^2} - \xi\theta = 0$, but this time with a pinned- (at $s = 0$) and a free-end (at $s = L$) as boundary conditions. This corresponds exactly to the classical case of a column buckling under its own weight, as considered in [10, 11], and the buckling occurs above $L_{p,c}^3 \approx 7.8 K_b / f_{\text{glide}}$.

With the parameter values provided in Table S1, estimated for *F. draycotensis* and other cyanobacteria, we predict that buckling occurs for filaments longer than $L_{r,c} \approx 146\mu$m for the compression scenario, and $L_{p,c}^3 \approx 92\mu$m for the pinned-head scenario. This is consistent with experimental results (see Figure 3D). Importantly, both values are lower than the threshold for twist instability and, in particular, plectoneme formation. This is consistent with the experimental observation that intermediate-length filaments can buckle but not form plectonemes (see Movie S1 and Figure 3D).

In addition, we showed that the load is not uniform and instead accumulates in the middle section of the filament that is compressed by the two sides moving against each other. As a result, when filaments buckle, we expect the buckling to be localised and not span the entire filament. This is indeed what we observe experimentally (see Movies S1 and S10-12). From our scaling arguments, the size of the buckled region is also related to the characteristic buckling length $l_0$.

#### Simulation results for a wide parameter range and effects of key parameters

We explore a broader range of parameter values and examine the dynamics of filaments under a decoordinated reversal, in particular, whether an instability occurs and what sets the shape of the filament during this instability. This approach enables a more systematic and controlled exploration of the parameter space than is feasible in experiments. However, simulating long, rigid filaments in the regime relevant to *F. draycotensis* is computationally expensive due to the numerical stiffness inherent in elasto-hydrodynamic simulations [12]. We therefore perform additional simulations using more flexible filaments (lower $K$ values).

The main free parameters in our system are the filament flexibility (relative bending modulus $K$), the length $L_f$, the bending-to-twisting moduli ratio $\Gamma$, and the adhesion strength $c_{adh}$. We explore their respective role in the filament dynamics and find that they are strongly coupled in setting the instability type (straight, buckled, plectoneme, knotted) and the filament shape during the reversal (see Figure 3E and S3 and Movie S8). Specific impacts of different parameters are discussed next.

***Bending-to-twisting moduli ratio.*** While the linear instability thresholds we computed analytically only depend on the bending stiffness $K_b$, we find that the ratio of twist-to-bend moduli $\Gamma$ has an important role in setting the shape of the filament during a reversal-induced-instability. In particular, for filaments that buckle but do not form plectonemes, we find that the bending mode changes with $\Gamma$ (Figure S3A). Filaments with low or moderate relative twisting moduli buckle in an even wave shape (mode 2 buckling). As $\Gamma$ increases, that is, the filament has a constant bending modulus but resists twisting more, the buckling mode changes to what seems to be an initial odd mode-3 buckling, and as the buckling progresses, the filament deformation occurs on one side only. The threshold between these buckling modes depends on filament stiffness and length (Figure 3A). We note that commonly observed experimental buckling events are of mode 2 type (see Movie S1).

***Adhesion.*** At low adhesion, we find that buckling occurs orthogonally to the surface, with the filament detaching away from it (see Figure S3B.i). Increasing the adhesion between the surface and the filament leads to buckling instead occurring on the surface. Very high adhesion strengths can also prevent plectoneme formation, as local detachment is necessary to form the head loop of the plectoneme (Figure S3B.ii). These model results, suggesting an optimal adhesion strength for plectoneme formation, are interesting given that plectonemes are readily observed on glass, while plectoneme formation on agar - where filament adhesion, via slime, is expected to be much stronger - is much more rare [7]. Moreover, we observed that the growth of *F. draycotensis* cultures on plastic surfaces - where higher adhesion forces might be possible - enhanced biofilm formation over granular macrostructure formation.

### Supplementary Tables

**Table 1** Model parameters and numerical parameters

| | | |
|---|---|---|
| Filament length | $L_f$ | $100 - 1000\mu$m |
| Bending modulus | $K = K_B/K_{B,0}$ | $K_{B,0} = 10^5$ nN.$\mu$m$^2$ [13, 14] (range $K = 0.01 - 1$) |
| Twisting modulus | $\Gamma = K_T/K_B$ | 1 (range $1 - 10$) |
| Gliding force density | $f_{\text{glide}}$ | 1 nN.$\mu$m$^{-1}$ [14] |
| Gliding torque density | $n_{\text{glide}}$ | 12.5 nN.$\mu$m$^{-1}$ [7] |
| Adhesion strength | $c_{\text{adh}}$ | $5f_{\text{glide}}$ |
| Environment viscosity | $\mu$ | 0.2nN.s.$\mu$m$^{-2}$ |
| Film thickness | $\delta$ | $1\mu$m |
| Discretisation size | d$s$ | 1 [$\mu$m] |
| Regularisation parameter | $\epsilon$ | 3 d$s$ |
| timestep | d$t$ | $10^{-5}[s]$ |
| Steric repulsion | $c_{\text{ste}}$ | $10^3$ |
| Adhesion range | $\delta_{\text{adh}}$ | 0.1 [$\mu$m] |

**Table 2** Laboratory and enrichment-culture genomes included in Figure S5. All were assembled from metagenomes of non-axenic laboratory cultures: *F. draycotensis* from PacBio HiFi reads of the laboratory culture [15], and all others from the enrichment cultures of Scarampi et al. [16]. Accessions are NCBI BioSamples under BioProject PRJNA1432497. Taxonomy was estimated with the GTDB release R232 [17, 18], with the corresponding order of Strunecký et al. [19] where available. Asterisks mark the two strains used in the particle-collection experiments. The *F. draycotensis* genome sequence and the alignment and trees underlying Figure S5 are provided in the Zenodo record of this study (see *Data and code availability*). All other genomes in Figure S5 were obtained from GTDB; the full list of 97 genomes, with accessions, classification, morphology and habitat, is given in Supplementary File S1.

| **Label in Figure S5 (genome)** | **Accession** | **GTDB order** | **GTDB family** | **Strunecky order** |
|---|---|---|---|---|
| *Fluctiforma draycotensis* * PacBio assembly | PRJNA1432497 | Cyanobacteriales | JAAUUE01 | – |
| *Pseudanabaena* sp. * OSS1_Bin_161 | PRJNA1432497 | Pseudanabaenales | Pseudanabaenaceae | Pseudanabaenales |
| JAAUUE01 sp. OSS9_Bin_80 | SAMN56435584 | Cyanobacteriales | JAAUUE01 | – |
| *Pseudanabaena biceps* SD3_Bin_60 | SAMN56435609 | Pseudanabaenales | Pseudanabaenaceae | Pseudanabaenales |
| *Cyanobium* sp. SD2_Bin_45 | SAMN56435557 | PCC-6307 | Cyanobiaceae | Synechococcales |
| WH-5701 sp. OSS6_Bin_56 | SAMN56435635 | PCC-6307 | Cyanobiaceae | – |
| *Geminocystis* sp. OSS12_Bin_158 | SAMN56435572 | Cyanobacteriales | Cyanobacteriaceae | Chroococcales |
| *Microcoleus* sp012035135 SD2_Bin_9 | SAMN56435600 | Cyanobacteriales | Microcoleaceae | Oscillatoriales |
| *Microcoleus* sp015207285 OSS9_Bin_52 | SAMN56435601 | Cyanobacteriales | Microcoleaceae | Oscillatoriales |
| *Nodosilinea* sp. OSS7_Bin_122 | SAMN56435602 | Phormidesmidales | Phormidesmidaceae | Nodosilineales |
| *Scytolyngbya* sp. OSS8_Bin_34 | SAMN56435619 | Leptolyngbyales | Leptolyngbyaceae | – |
| *Calothrix* sp. NONANO_MAG_00047 | PRJNA1432497 | Cyanobacteriales | Nostocaceae | Nostocales |

### Supplementary Figures

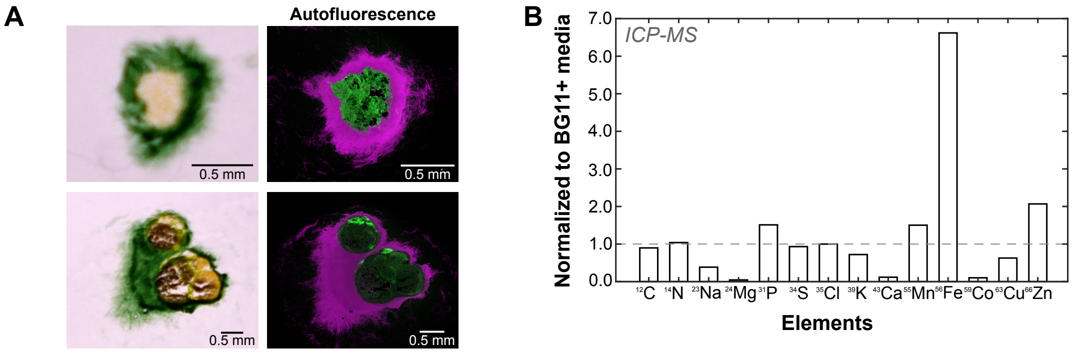


C

| Elemental Composition (atomic %) | | | | | | | |
|---|---|---|---|---|---|---|---|
| Fe | Mn | O | C | N | Ca | P | Mg |
| 1.63 | 0.38 | 38.24 | 49.21 | 2.69 | 3.20 | 3.63 | 1.02 |

**Fig. 1 Granule structure and iron analysis.** (a) Example cryosliced images of macrostructures. Thin slices of the granules show the yellow/brown core and surrounding cyanobacteria, compared with comparative autofluorescence obtained on a Zeiss 880 confocal microscope (green: ex.: 488 nm, em.: 540-580 nm; magenta: ex. 561 nm, em.: 640-690 nm). Magenta regions show the autofluorescence of the cyanobacterial chlorophyll. (B) ICP-MS analysis of the extracted yellow core. Elemental composition of the sample, normalised to the concentration found in an equivalent sample mass of BG11+. (C) XPS analysis of the extracted yellow core, showing elemental composition as atomic percentage. Note that whilst the granule may be enriched with iron compared to the media conditions, the yellow material is predominantly of an organic nature (49.21% carbon), with 1.63% of the composition attributable to iron.

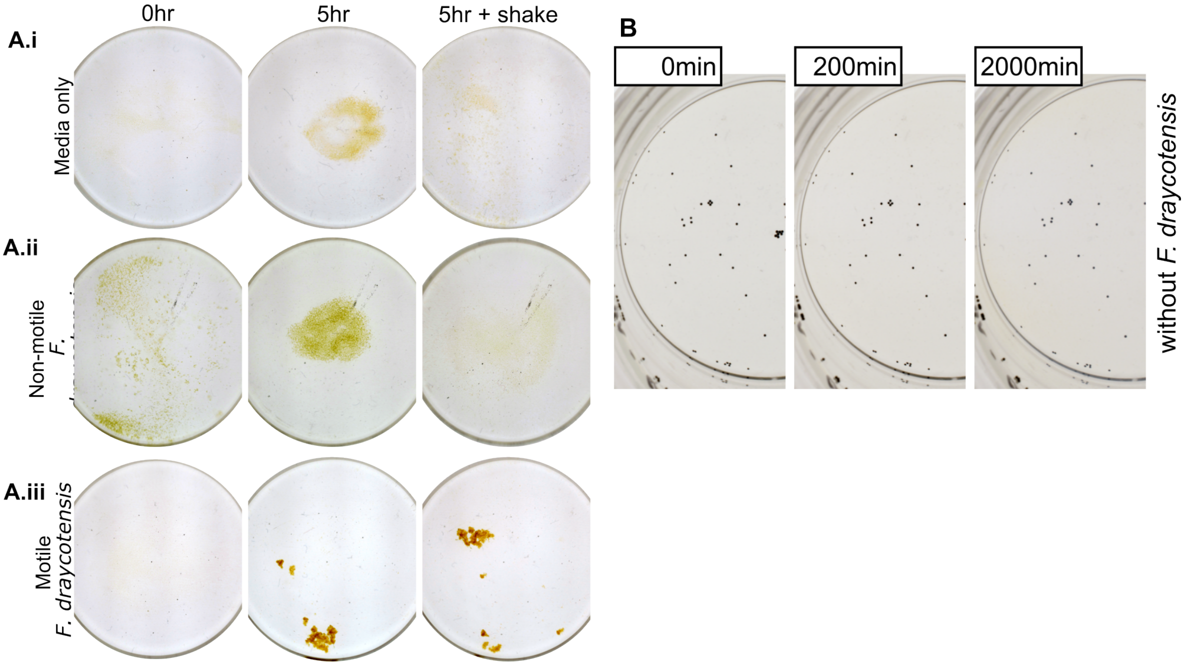


**Fig. 2 Comparison of motile cyanobacteria, non-motile cyanobacteria, and no-cyanobacteria cultures and media-only control** (A) Particle collection assay in a glass beaker for (i) sterile BG11+ media, (ii) a community culture containing a mutant strain of *F. draycotensis* that does not exhibit motility, and (iii) the normal *F. draycotensis* community culture. Images show the beakers at initiation of experiment, and their appearance after 5 hours: firstly undisturbed and then following an additional agitation by hand (swirling and tapping beaker on the bench) to redistribute the particles. (B) Particle collection assay with a derived community lacking *F. draycotensis*. Note that these experiments were initiated containing black polystyrene beads to track any collection(see *SM*).

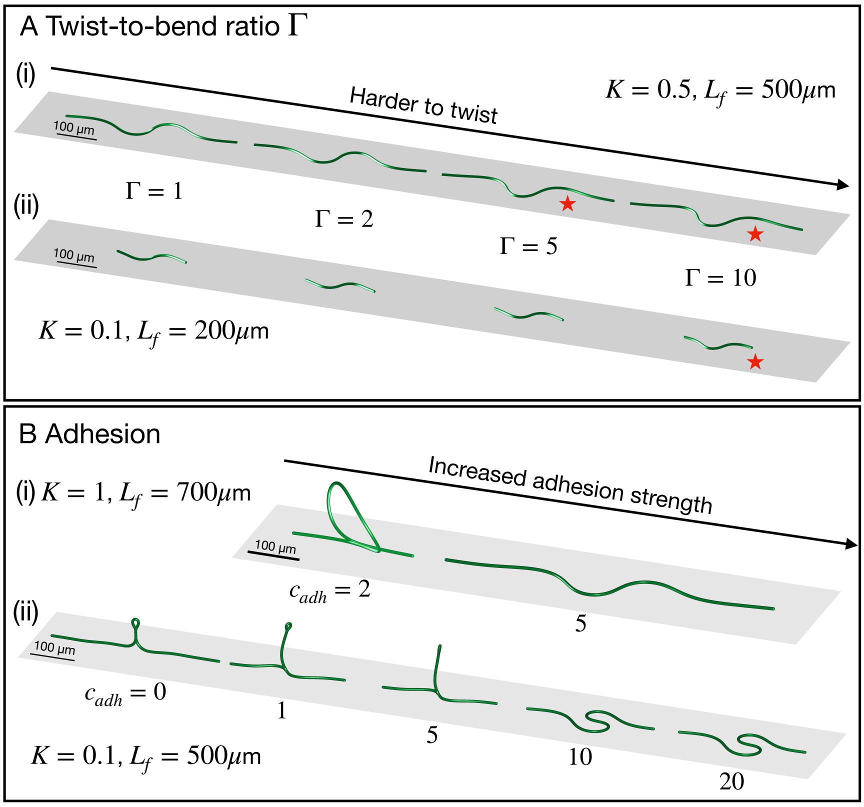


**Fig. 3 Mechanical properties and adhesion set the shape of simulated buckling filaments.** (A) Shape of buckled flexible filaments after 120 s when increasing the relative twist bending modulus $K_t = \Gamma K$ for a given bending stiffness. At higher $\Gamma$, we observe a transition from symmetric buckling (wave shape) to a one-sided buckling resembling experiments, as marked by red stars. (B) Increasing the strength of the surface adhesion leads to buckling in the plane of the surface instead of orthogonally detached from it (top panel) and can prevent plectoneme formation as the loop at the plectoneme head doesn't form (bottom panel).

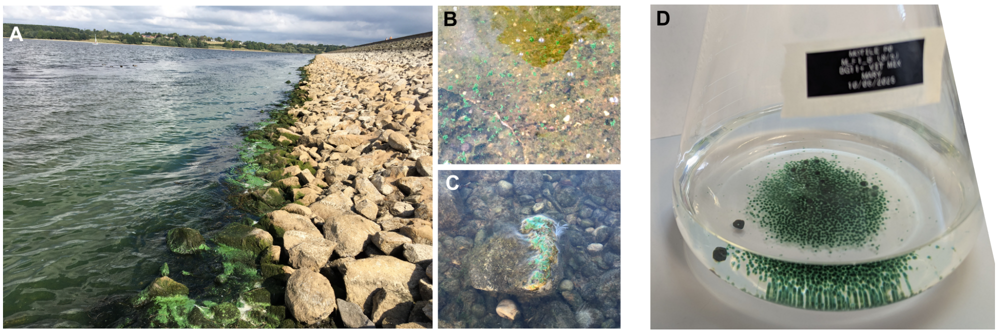


**Fig. 4** **Cyanobacteria in alternative growing conditions.** (A-C) Photographs showing growing conditions and morphologies of cyanobacteria in the original lake environment. (A) High densities of cyanobacteria at the shoreline. (B) Macroscopic green granules (likely to be a species of cyanobacteria other than *F. draycotensis*) in the water column. (C) Cyanobacterial mat on a partially submerged rock. (D) Growing a culture under continuous rotational flow promotes the formation of granular macrostructures in our laboratory community.

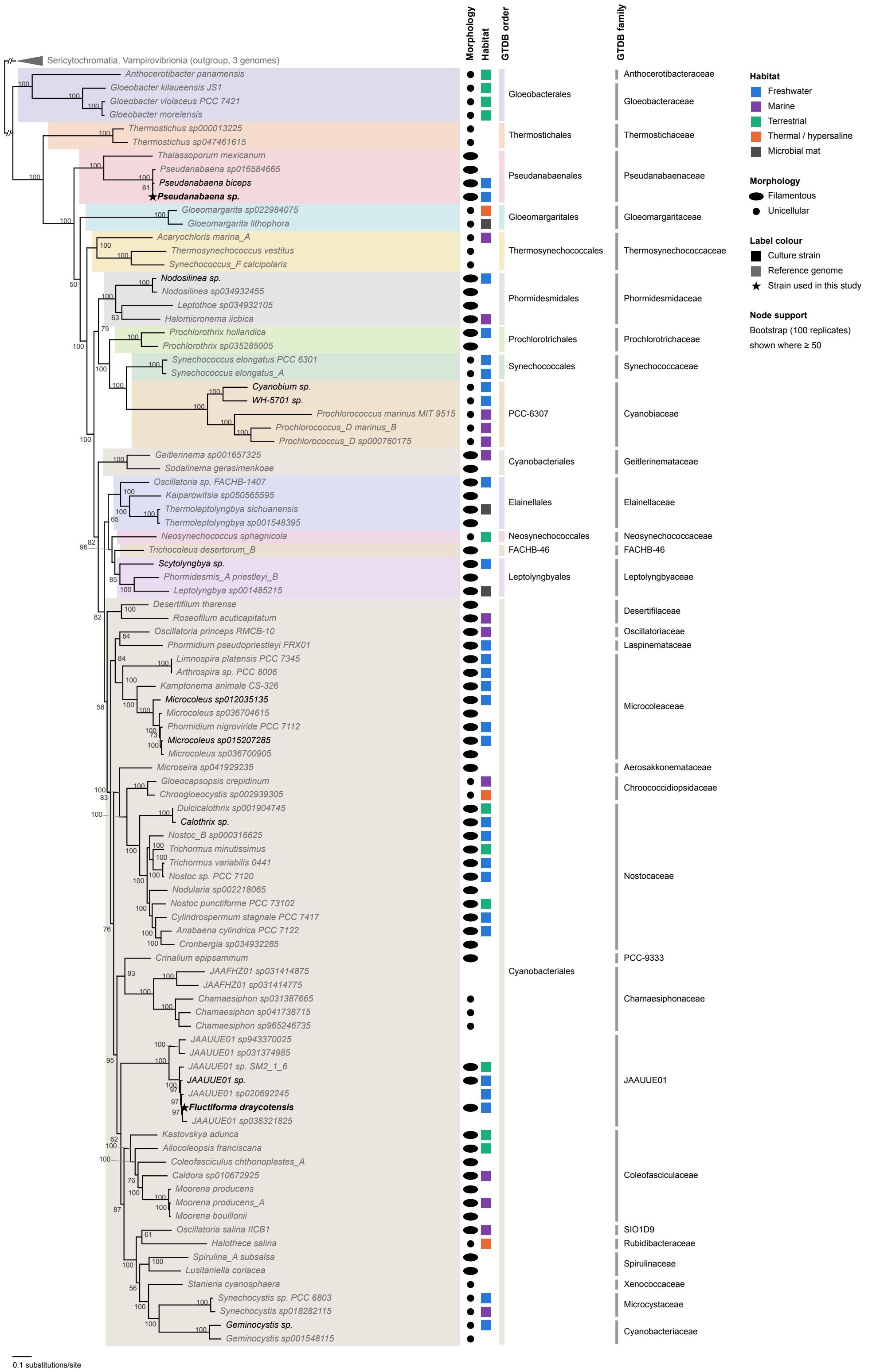


**Fig. 5 Phylogenomic placement of *F. draycotensis* and *Pseudanabaena* sp.** Maximum-likelihood tree of 94 cyanobacterial genomes inferred with IQ-TREE 3 from 67 concatenated single-copy marker proteins (12,293 amino-acid positions; see *Methods*), rooted on three non-photosynthetic Cyanobacteriota (Sericytochromatia and Vampirovibrionia), shown collapsed at the top; the two root branches (//) are not drawn to scale. Shaded blocks and the columns on the right give the order and family of each genome in the Genome Taxonomy Database (GTDB, release R232) [17, 18]. The order of each genome under Strunecký et al. [19] is listed in Supplementary File S1. Numbers at nodes are non-parametric bootstrap percentages (100 replicates). Symbols at the tips give morphology (filamentous or unicellular) and habitat (thermal and hypersaline sites are combined; the habitat of each genome is listed in Supplementary File S1); blank cells mean no information was available. Label colours distinguish culture strains (black) from reference genomes (grey), and stars mark the two strains used in this study. Scale bar, 0.1 amino-acid substitutions per site.

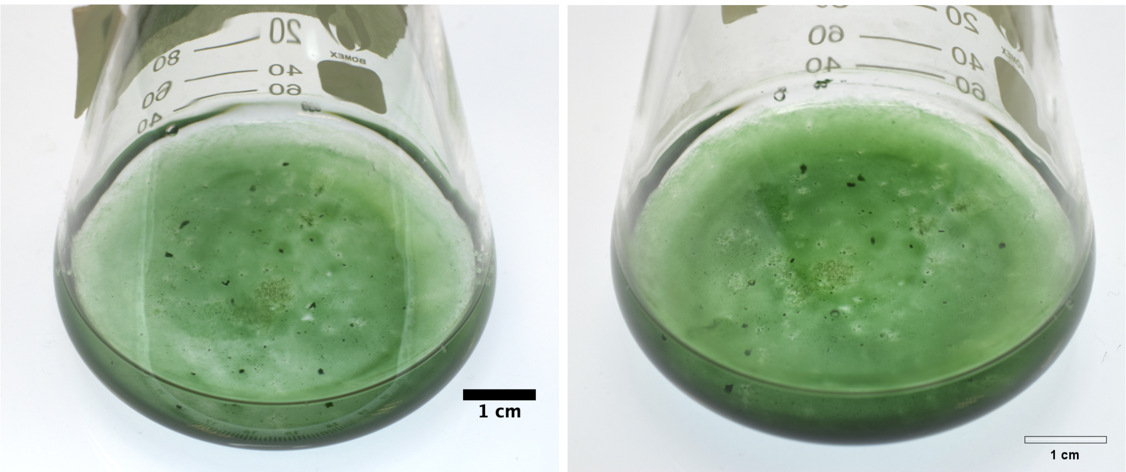


**Fig. 6 A *Pseudanabaena* culture at 16 and 23 days of growth.** No granular macrostructures are visible: instead, the cyanobacteria form a biofilm that adheres strongly to the surface of the beaker. Some denser regions are observed, but they are not granular as seen in the *F. draycotensis* cultures.

#### Additional Supplementary files

- *poonEtAl_supplementaryFile1.tsv* Supplementary File S1. The full list of 97 genomes used in the phylogenetic tree shown in Figure S5 (those obtained from GTDB and from laboratory cultures). The list includes accession number, GTDB and Strunecky classification for each genome, along with the morphology and habitat of the species, where known.
- *poonEtAl_MovS1.mov* Movie S1.Exemplar movie showing gliding motility in a *F. draycotensis* population, displaying characteristic back-and-forth trajectories.
- *poonEtAl_MovS2.mov* - Movie S2. Macroscale particle collection and macrostructure formation. Exemplar movie showing formation of yellow particles and their collection by *F. draycotensis* filament bundles, and forming of macrostructures. Still images from this movie are shown in Figure 1C.
- *poonEtAl_MovS3.mov* - Movie S3. **Macroscale particle collection by filament bundles.** Exemplar movie showing collection of particles by *F. draycotensis* filament bundles, and coalescence of structures. Part of this movie is shown as a static image in Figure 1D.
- *poonEtAl_MovS4.mov* - Movie S4. **Collection of macroscopic beads.** Exemplar movie showing collection of macroscopic beads added to the culture media by *F. draycotensis* filament bundles, and coalescence of structures.
- *poonEtAl_MovS5.mov* - Movie S5. **Microscale particle collection by filaments.** Exemplar movie showing collection of microscopic particles by *F. draycotensis* filaments. Part of this movie is shown as a static image in Figure 2C.
- *poonEtAl_MovS6.mov* - Movie S6. **Microscale particle collection by a single filament.** Exemplar movie showing a single twisting *F. draycotensis* filament, collecting microscopic particles.
- *poonEtAl_MovS7.mov* - Movie S7. **Simulation of plectoneme formation in a reversing filament.** Animated version of Figure 3C.ii.
- *poonEtAl_MovS8.mov* - Movie S8. **Simulation of phase space: dynamics of revering filaments of different length and flexibility.** Animated version of Figure 3E. Rigid and short filaments stay straight or buckle on the surface, while longer, more flexible ones can form plectonemes. Very flexible filaments end up knotting, with more loops forming along the filament.
- *poonEtAl_MovS9.mov* - Movie S9. **Simulation of two filaments gliding over each other: separation and entanglement.** Animated version of Figure 3F. In pairs of filaments ($K = 0.1$), shorter filaments (in the straight or buckling regime) end up separating, while longer, plectoneme-forming filaments entangle. The green filament is gliding forward, while the orange one is reversing.
- *poonEtAl_MovS10.mov* - Movie S10. **Two filaments meeting without entanglement.** Exemplar movie showing two *F. draycotensis* filaments meeting, and one buckling, but the two not entangling.
- *poonEtAl_MovS11.mov* - Movie S11. **Two filaments meeting and entangling.** Exemplar movie showing two *F. draycotensis* filaments meeting, one forming a plectoneme and entangling the other.
- *poonEtAl_MovS12.mov* - Movie S12. **Multiple filaments meeting and entangling.** Exemplar movie showing several *F. draycotensis* filaments, one forming a plectoneme and initiating a larger entanglement of multiple filaments.
- *poonEtAl_MovS13.mov* - Movie S13. **Gliding motility in *Pseudanabaena*.** Exemplar movie showing *Pseudanabaena* filaments gliding on a glass surface.
